# Brain Regional Gene Expression Network Analysis Identifies Unique Interactions Between Chronic Ethanol Exposure and Consumption

**DOI:** 10.1101/688267

**Authors:** M.L. Smith, M.F. Lopez, A.R. Wolen, H.C. Becker, M.F. Miles

**Author notes:** Corresponding authors: Michael F. Miles, Howard C. Becker.

## Abstract

Progressive increases in ethanol consumption is a hallmark of alcohol use disorder (AUD). Persistent changes in brain gene expression are hypothesized to underlie the altered neural signaling producing abusive consumption in AUD. To identify brain regional gene expression networks contributing to progressive ethanol consumption, we performed microarray and scale-free network analysis of expression responses in a C57BL/6J mouse model utilizing chronic intermittent ethanol by vapor chamber (CIE) in combination with limited access oral ethanol consumption. This model has previously been shown to produce long-lasting increased ethanol consumption, particularly when combining oral ethanol access with repeated cycles of intermittent vapor exposure. The interaction of CIE and oral consumption was studied by expression profiling and network analysis in medial prefrontal cortex, nucleus accumbens, hippocampus, bed nucleus of the stria terminalis, and central nucleus of the amygdala. Brain region expression networks were analyzed for ethanol-responsive gene expression, correlation with ethanol consumption and functional content using extensive bioinformatics studies. In all brain-regions studied the largest number of changes in gene expression were seen when comparing ethanol naïve mice to those exposed to CIE and drinking. In the prefrontal cortex, however, unique patterns of gene expression were seen compared to other brain-regions. Network analysis identified modules of co-expressed genes in all brain regions. The prefrontal cortex and nucleus accumbens showed the greatest number of modules with significant correlation to drinking behavior. Across brain-regions, however, many modules with strong correlations to drinking, both baseline intake and amount consumed after CIE, showed functional enrichment for synaptic transmission and synaptic plasticity.

## Introduction

Alcohol use disorder (AUD) is a highly significant public health issue. The condition contributes to over 60 types of diseases, and is responsible for over 2 million deaths worldwide every year [1, 2]. A hallmark of AUD is progressive, abusive ethanol consumption over time. This increase in ethanol consumption is thought to be due to neurobiological adaptations induced by ethanol itself, and the repeated occurrence of ethanol withdrawal [3]. Previous studies in humans and animal models of chronic alcohol exposure have led to the hypothesis that changes in gene expression are a major molecular mechanism contributing to physiological and behavioral alterations accompanying AUD [4–7].

Technologies such as microarrays have allowed for the study of the genome-wide effects of ethanol exposure on mRNA expression [4], and scale-free network analysis provides a means to organize transcriptome data into networks of co-expressed genes representing functional pathways [8–12]. Further, gene-phenotype correlations allow for the identification of both individual genes and gene networks associated with dependent variables such as ethanol consumption. Using such approaches it may be possible to identify molecular network functions contributing to increased drinking behavior seen with chronic ethanol exposure, and to pinpoint candidate genes whose expression correlates with consumption; thus identifying new potential therapeutic targets for the treatment of AUD.

Recent studies provide substantial predictive validation of new animal experimental models for the discovery of therapeutic targets in the treatment of AUD [13–16]. Chronic intermittent ethanol vapor exposure (CIE) in rodents is one such model, providing long-term intermittent intoxicating ethanol exposure. As a part of this paradigm, mice or rats experience repeated cycles of high blood ethanol levels provided by vapor exposure followed by withdrawal, similar to behavioral patterns seen in alcoholics [17]. The CIE by vapor chamber model has been shown to cause neurochemical and structural changes at the synapse and increases in ethanol consumption. Our laboratories have also previously identified complex brain region-specific temporal patterns of gene expression changes upon withdrawal from CIE [11, 18]. In mouse models, providing limited access 2-bottle choice ethanol consumption in between cycles of ethanol vapor exposure has been shown to more rapidly and significantly increase ethanol consumption [19]. The molecular mechanisms underlying this combined action of voluntary oral ethanol consumption and cycles of high dose ethanol withdrawal on ethanol consumption are unknown but could provide important directions for possible intervention in the progression from social to abuse drinking [19, 20]. Previous studies of gene expression with CIE in C57BL/6J mice have focused on differential gene expression during early withdrawal [21], or on RNA networks during ethanol exposure and withdrawal associated with cell type-specific gene expression [18]. This current study explores the relationship between high-dose ethanol vapor exposure, intermittent drinking, and withdrawal in an attempt to identify mechanisms by which this model leads to progressive increases in ethanol intake.

This manuscript presents a detailed analysis of gene expression network-level changes caused by CIE exposure with or without intermittent oral ethanol consumption, across multiple brain-regions using Weighted Gene Correlated Network Analysis (WGCNA) [22]. The brain-regions studied have been associated in numerous studies with the development of AUD [15, 23, 24]. By combining statistical analysis for genes regulated by ethanol consumption, CIE, or the combination, we identify brain-region selective expression networks responding to particular ethanol exposure models. Specifically we show that prefrontal cortex (PFC) and bed nucleus of the stria terminals (BNST) showed prominent responses to CIE and drinking, but the nucleus accumbens (NAC) and hippocampus (HPC) were primarily responsive to high-dose ethanol vapor exposure alone, while gene expression in the central nucleus of the amygdala (CeA) may be particularly altered by ethanol withdrawal. Furthermore, we identified expression networks that correlated with increased ethanol consumption caused by cycles of CIE and drinking, suggesting mechanistic relationships. We also demonstrate that some of the most strongly correlated genes are those related to synaptic transmission and synaptic plasticity. Together, our findings contribute substantial new knowledge to our understanding of brain regional gene network adaptations contributing to brain plasticity during various stages of AUD.

## Materials and Methods

### Animals

Adult male C57BL/6J mice were purchased from Jackson Laboratories (Bar Harbor, ME, USA) at 10 weeks of age. Mice were kept under a 12-hour light/dark cycle and given free access to water and standard rodent chow (Harland, Teklad, Madison, WI). Mice were kept on corncob bedding (#7092a and #7902.25 Harland, Teklad, Madison, WI). All studies were conducted in an AALAC-accredited animal facility, and approved by the Institutional Animal Care and Use Committee at Medical University of South Carolina (MUSC). All experimental and animal care procedures met guidelines outlined in the NIH Guide for the Care and Use of Laboratory Animals.

### Chronic Intermittent Ethanol (CIE)

Studies were designed to determine genomic responses and interactions between two different ethanol exposure models: intermittent cycles of ethanol vapor exposure in inhalation chambers (CIE), and oral consumption of 15% (v/v) ethanol in a limited access (2 h/session) paradigm. Mice were divided into 4 treatment groups: the CIE-Drinking group received inhaled ethanol in the vapor chambers followed by 2-bottle choice ethanol drinking in between vapor exposure cycles; the Air-Drinking group received only air in the vapor chambers, but had 2-bottle choice ethanol drinking between CIE cycles; the CIE-NonDrinking group received inhaled ethanol in the vapor chambers but only water access in between CIE cycles; and the Air-NonDrinking group remained ethanol naïve with air exposure in vapor chambers and only water consumption between CIE cycles. Following a 2-week acclimation period, mice in the CIE-Drinking and Air-Drinking groups underwent 6-weeks of 2-bottle choice drinking to establish baseline drinking levels. Ethanol and water intake for each individual mouse was measured daily. Following 6-weeks of baseline drinking, mice were placed in Plexiglass inhalation chambers (60×36×60 cm) 16 hours/day for 4 days. Ethanol was volatilized with an air stone submerged in 95% ethanol. Vapor chamber ethanol concentrations were monitored daily and air flow was adjusted to ethanol concentrations within 10-13 mg/l air. This ethanol vapor concentration has been shown to yield stable blood ethanol concentrations (175-225 mg/dL) in C57BL/6J mice [25]. Before each vapor chamber session, intoxication was initiated in the CIE group by administration of 1.6 g/kg ethanol and 1 mmol/kg pyrazole intraperitoneally (i.p.) at a volume of 0.02 ml/g body weight. Pyrazole is an alcohol dehydrogenase inhibitor used to stabilize blood ethanol concentrations. All mice received the same number and timing of pyrazole injections prior to final removal from the inhalation chambers with control mice receiving saline and pyrazole (i.p.), also at a volume of 0.02 ml/g body weight, prior to being placed into control vapor chambers. Control vapor chambers delivered only air without ethanol vapor. After 4 days in the inhalation chambers, mice underwent a 72-hour period of total abstinence from ethanol. Following the abstinence period, mice in the CIE-Drinking and Air-Drinking groups were given 2-bottle choice drinking for 2 hours per day for 5 days. A total of 4 cycles of CIE-abstinence-drinking were performed. After the end of the 4^th^ cycle mice were sacrificed on the 5^th^ drinking day before receiving ethanol/water access on that day at the time they received 2-bottle choice drinking all previous drinking days (Figure 1).

**Figure 1:**
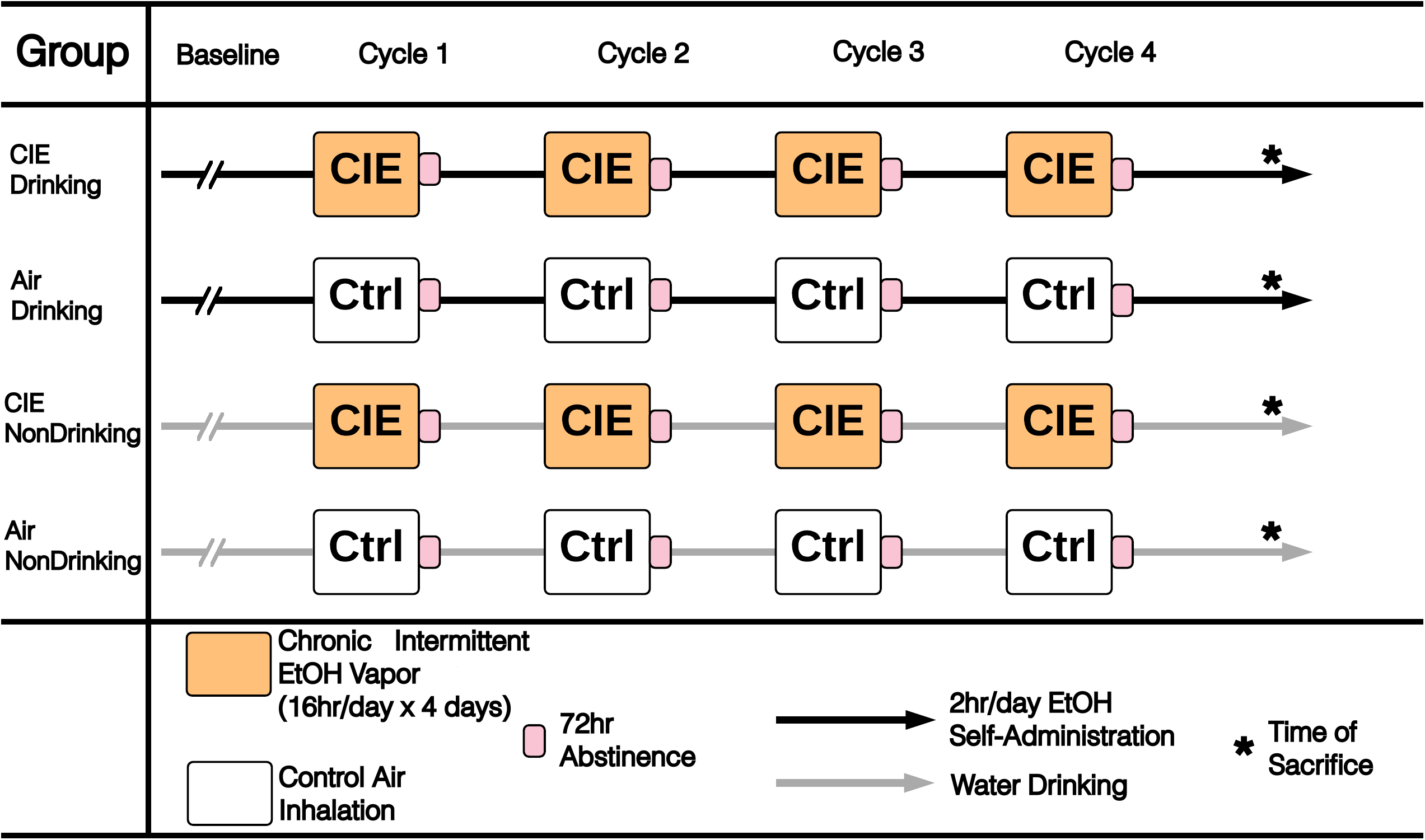
Schematic representation of CIE and drinking experimental design.

### Tissue Harvesting, RNA Isolation, and Microarray Hybridization

Mice were sacrificed by decapitation, brains were immediately removed from the skull, and brain-regions dissected as previously described [21]. Tissues were stored at −80°C until RNA isolation. Total RNA was extracted using the RNeasy Mini Kit (Qiagen Valencia, CA). Affymetrix GeneChip® Mouse Genome 430, type 2 arrays were used to measure gene expression. Sample preparation, hybridization, and array scanning were performed at the MUSC ProteoGenomics Core Facility according to procedures optimized by Affymetrix (Santa Clara, CA, USA). Each brain-region was processed separately with treatment groups randomized to minimize batch effects. Array data was stored in CEL file format, and sent to Virginia Commonwealth University (VCU) for analysis.

### Microarray Analysis

Affymetrix GeneChip® Mouse Genome 430, type 2 arrays were analyzed with The R Project for Statistical Computing (http://www.r-project.org/). Microarray quality was assessed by RNA degradation, average background, percent present probesets, and multi-dimensional scale plots (first principal component by second principal component). Arrays showing low quality measures, or that appeared to be outliers, were removed from the dataset. Background correction using Robust Multi-array Average (RMA) and quantile normalization was performed using the affy package for R [26, 27]. Each brain-region was normalized separately. ComBat by RNA hybridization batch was used to correct for any batch effects present in the data [28].

### CIE and Drinking Responsive Genes

Statistical analysis to identify significantly regulated genes was performed using the limma package for R [29]. Two factor LIMMA looking at treatment and drinking, as well as interaction, was used for initial analysis. However, we also ran LIMMA with each treatment group as an independent variable. This was done based on the fact that, over the course of the study, each group received a different overall dose of ethanol, number of, and duration of exposure. Each possible comparison between the 4 treatment groups was performed leading to 6 total comparisons labeled 1 through 6. Overall significance was also measured by ANOVA. Multiple testing was adjusted using the Benjamini and Hochberg false discovery rate method (FDR) [30]. False discovery rates equal to or less than 0.01 were considered significant.

### Statistical Analysis of 2-Bottle Choice Drinking

Average ethanol intake (g/kg) was calculated across 5 drinking days of each week during the baseline-drinking period. During the testing cycles, mice also drank for 5 days; therefore average drinking across these 5 days was calculated to represent drinking during each CIE cycle. Differences in drinking were determined by Two Way ANOVA with Repeated Measures using SigmaPlot 12.0 (Systat Software, San Jose, CA, USA).

### Weighted Gene Correlated Network Analysis

Weighted Gene Correlated Network Analysis (WGCNA) was used to perform scale-free network topology analysis of microarrays [22]. Such scale-free network approaches have been used previously to identify biological pathways influenced by ethanol exposure in mice [11, 12]. WGCNA was performed on each brain-region separately using the WGCNA package for R [31]. Overall significance by one-way ANOVA comparing all groups (FDR equal to or less than 0.01) was used to select probesets to be included in network analysis. A probeset found to be significant by ANOVA in any brain region was included to generate the overall probeset list used for WGCNA across all brain regions. Standard WGCNA parameters were used for analysis with the exceptions of soft-thresholding power and deep split. Appropriate soft-thresholding powers were selected using previously described methods [31]. A soft-thresholding power of 6 was used for all brain-regions except the PFC for which a soft-thresholding power of 8 was used. WGCNA was performed with deep-split values of 0-3. Deep-split value was selected by a multi-dimensional scaling (MDS) plot, which displayed first and second principal components. Deep-split values were to minimize module overlap on the MDS plot. Deep-split values of 3 were chosen for the PFC, NAC, and CeA. For the HPC a deep-split of 2 was chosen, and a deep-split of 0 for the BNST. Modules were validated based a permutation procedure outlined by Iancu et al. [32]. Briefly, the average topological overlap of probesets assigned to each module was compared to the average topological overlap of 100 bootstrapped modules comprised of randomly sampled probesets. Z-scores of average topological overlap between probesets assigned to the module, and modules comprised of random probesets were used to calculate p-values and false discovery rates (FDR). Modules with FDR values ≤ 0.2 were considered validated.

### WGCNA-Drinking Correlation

Modules identified by WGCNA were related to drinking data by Spearman Rank correlation using the module eigengene as previously described [33, 34]. Individual probesets were also correlated to drinking data with the Spearman Rank method. These correlations were then used to identify modules enriched in genes whose expression showed systemic relationships with drinking behavior across 4 cycles of CIE with 2-bottle choice drinking.

### Bioinformatics

Modules identified by WGCNA were examined for function using publicly available bioinformatics resources. The Functional Annotation Chart tool from DAVID (http://david.abcc.ncifcrf.gov/) [35] was used to identify biological pathways highly represented by genes grouped into each module. Gene Ontology terms were then summarized by semantic similarity using REVIGO (revigo.irb.hr/). GeneMANIA (http://www.genemania.org) was also used for functional analysis through use of GO process constituent genes as query lists. GeneMANIA mines public database and publication data to identify known associations between genes and their protein products. Co-expression modules identified in this dataset were also compared to those identified in corresponding brain-regions in a previously published study from our laboratory of the time-dependent effects of multiple cycles of CIE by vapor chamber [11]. This comparison was performed using WGCNA’s userListEnrichment() function, utilizing hypergeometric overlap to determine significance of enrichment [31]. Hypergeometric overlap p-values were adjusted for multiple testing using false discovery rates [30]. Module overlaps were considered significant at a FDR ≤ 0.05. Since all brain-regions in this study used RNA from whole tissue samples, modules were also examined for enrichment for genes expressed in specialized cell-types [36] found in brain (neurons, astrocytes, and oligodendrocytes) to determine whether any identified modules represented specific cell-type gene expression changes within a brain-region [37]. The userListEnrichment() function was also used for cell-type enrichment analysis, with Bonferroni corrected p-values ≤ 0.05 considered significant.

### Module Disruption

Changes in network structure were measured based on the module disruption method outlined by Iancu *et al.* [38]. This method was adapted from the module preservation method [39], which examines module statistics across randomly selected network nodes (genes). The module disruption method looked at a set of bootstrap networks (n=200) generated by randomly selecting a subset of samples without regard to treatment group. Connectivity statistics as described by WGCNA [39] were then generated for each random network. The average correlation of each network’s intramodular connectivity (kIM) and total network connectivity (kME) to that for all other randomly generated networks was calculated. These values were then compared to the correlation of intramodular connectivity (cor.kIM) and total connectivity (cor.kME) between two treatment groups. For the purposes of this study, we compared the CIE Drinking group to the Air NonDrinking group. Difference in correlation between treatment groups, and all bootstrap networks were quantified using a Z score:

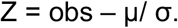

obs = Correlation between network statistic between treatment groups.

μ = Average correlation between network statistic for comparisons of 200 random networks.

σ = Standard deviation of correlation between network statistic for comparisons of 200 random networks.

In accordance with Iancu *et al.* [38], modules with Z scores less than −2 were considered significantly disrupted (see Suppl. Table 12).

## Results

### 2-Bottle Choice Drinking

Consistent with previous behavioral studies of CIE combined with ethanol consumption [19], Two Way ANOVA with Repeated Measures revealed significant differences in ethanol intake (p-value ≤ 0.05) between the CIE-Drinking and Air-Drinking groups after the first, third, and fourth vapor chamber session. After the second vapor chamber cycle, the CIE-Drinking group decreased ethanol intake compared to the first vapor chamber cycle, therefore, at this time-point, there was no significant difference in amount of ethanol consumed between CIE-Drinking and Air-Drinking groups. However, after the third and fourth vapor chamber sessions, the CIE-Drinking group drank significantly more ethanol than the Air-Drinking group (Figure 2, Suppl. Table 1). Interestingly, both the CIE-Drinking and Air-Drinking groups drank significantly more, compared to baseline, after only one session in the vapor chamber (Figure 2, Suppl. Table 1). This suggests that exposure to the air inhalation chambers may affect ethanol consumption. However, animals exposed to ethanol vapor during inhalation chamber sessions consumed significantly more ethanol, indicating that prolonged exposure to intoxicating levels of ethanol is the major driver of changes in drinking behavior.

**Figure 2:**
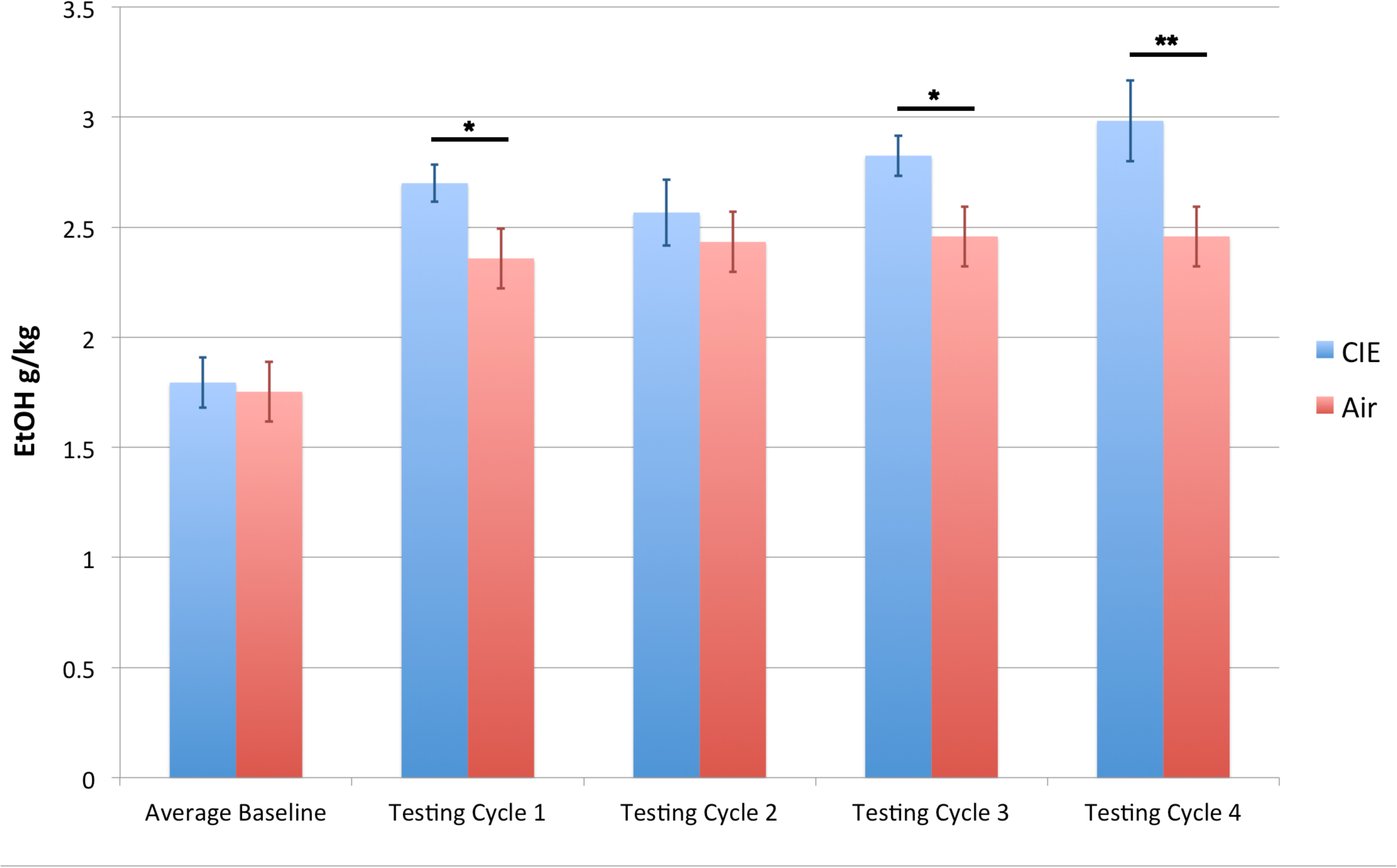
Ethanol intake in CIE and air control mice. Statistical difference in ethanol intake measured by two-way ANOVA with repeated measures, * p-value ≤ 0.05, ** p-value ≤ 0.01.

### Gene Expression with CIE and Drinking

To define molecular mechanisms contributing to the actions of chronic ethanol vapor exposure and intercurrent oral ethanol intake on escalating ethanol consumption seen in Figure 2, we performed extensive microarray studies across brain regions implicated in CIE through our prior genomic studies [11, 21]. Statistical analysis of microarray data with LIMMA found more significant differences in gene expression when each treatment group was treated as an independent group (Table 1, Suppl. Table 3). Significant differences in gene expression were found between each of the four treatment groups in the PFC, suggesting prominent treatment-specific responses in that brain region. Other brain regions, however, showed very different patterns of differential gene expression. In the NAC, HPC, BNST, and CeA, significant differences in gene expression were seen only seen with comparisons 1 (CIE-Drinking vs. Air-Drinking), 3 (CIE-Drinking vs. CIE-NonDrinking), and 4 (CIE-Drinking vs. Air-NonDrinking) (Table 1, Suppl. Table 2). Examining overlap between these comparisons revealed that a substantial number of genes were significant across all 3 comparisons, or between any combination of 2 comparisons in the PFC, BNST, and CeA. However, in the NAC and HPC, the majority of overlap was between CIE-Drinking vs. Air-NonDrinking and CIE-Drinking vs. Air-Drinking (Figure 3). Across NAC, HPC, BNST and CeA, the largest number of differentially expressed genes was seen between the CIE-Drinking group and the ethanol naïve Air-NonDrinking group (Table 1, comparison 4). These four regions, however, did show significant differential gene expression in comparison 1 (CIE-Drinking vs. Air-Drinking), and comparison 3 (CIE-Drinking vs. CIE-NonDrinking) (Table 1). This finding indicates an interaction between prolonged exposure to inhaled ethanol and voluntary intermittent drinking. Unique to the PFC, large expression differences were seen across all comparisons but comparison 4 (CIE-Drinking vs. Air-NonDrinking) had the smallest number of changes, in contrast to other brain regions (Table 1, Suppl. Table 2).

**Table 1:** Significantly differentially expressed probsets and genes between all comparisons of 4 treatment groups. Cells contain number of significant probesets and number of genes in parenthesis. Significant differential expression: LIMMA group comparisons, FDR ≤ 0.01.

| Table 1 | Comparison1 | Comparison2 | Comparison3 | Comparison4 | Comparison5 | Comparison6 |
| --- | --- | --- | --- | --- | --- | --- |
|  | CIE-Drinking<br>vs. Air-Drinking | CIE-NonDrinking<br>vs. Air-NonDrinking | CIE-Drinking<br>vs. CIE-NonDrinking | CIE-Drinking<br>vs. Air-NonDrinking | CIE-NonDrinking<br>vs. Air-Drinking | Air-Drinking<br>vs. Air-NonDrinking |
| PFC | 840<br>(775) | 569<br>(527) | 843<br>(764) | 325<br>(306) | 759<br>(705) | 629<br>(573) |
| NAC | 127<br>(122) | 0<br>(0) | 9<br>(9) | 1219<br>(1069) | 1<br>(1) | 0<br>(0) |
| HPC | 549<br>(502) | 0<br>(0) | 37<br>(37) | 1615<br>(1395) | 0<br>(0) | 0<br>(0) |
| BNST | 419<br>(391) | 0<br>(0) | 178<br>(168) | 543<br>(508) | 0<br>(0) | 0<br>(0) |
| CeA | 78<br>(76) | 0<br>(0) | 721<br>(641) | 818<br>(726) | 0<br>(0) | 0<br>(0) |

**Figure 3:**
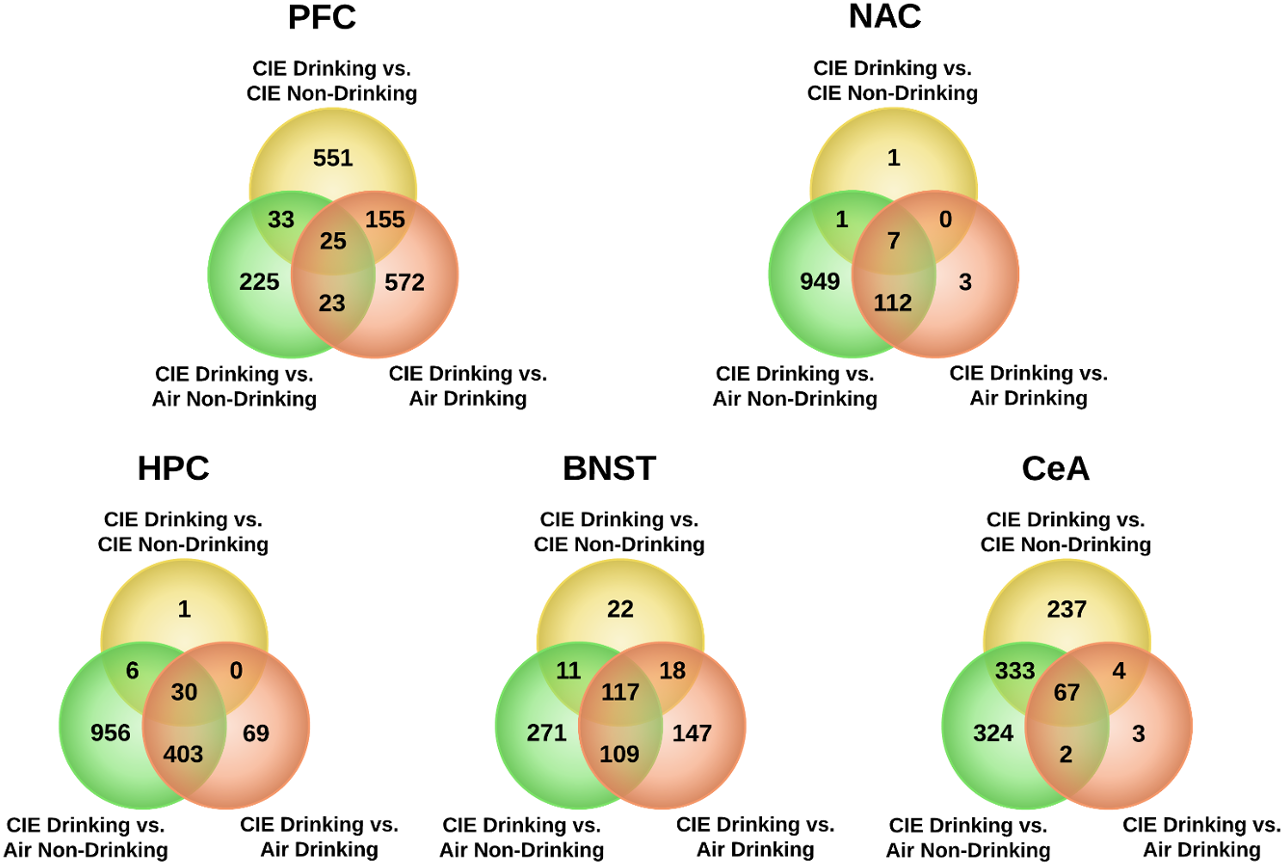
Overlap between 3 treatment/drinking group comparisons in all brain-regions. Overlap venn Diagrams of differentially expressed genes in PFC, NAC, HPC, BNST, and CeA between comparison of the CIE Drinking group and CIE control (CIE Non-Drinking), drinking control (Air Drinking), and ethanol naïve control (Air Non-Drinking). Significant differential expression: LIMMA FDR ≤ 0.01.

### Weighted Gene Correlated Network Analysis

To identify networks of coordinately regulated genes that might point to specific biological functions, we performed WGCNA analysis independently across all brain regions. To limit the WGCNA input to those genes showing some expression response to ethanol, we combined LIMMA-positive gene lists across all brain regions as described in Methods and our prior studies [11]. WGCNA identified modules of co-expressed genes in all brain-regions. Module sizes varied from over 3000 probesets to less than 35 (Table 2). Module validation using topological overlap showed that most modules identified withstood permutation of constituent genes, as indicated by Z-score false discovery rates ≤ 0.2 (Suppl. Table 11). One module in the CeA, greenyellow, did not show a significant false discovery rate, indicating that module may be the result of spurious associations. Additionally, in each brain region, the FDR of topological overlap Z-scores for the grey modules was 1 as expected, since WGCNA groups all genes which do not show significant topological overlap with any other module into the grey module. When WGCNA modules were interrogated by over-representation analysis for LIMMA-positive genes from various treatment comparisons across brain regions, PFC showed the largest extent of enrichment for LIMMA-positive genes across modules and these were generally distributed across multiple treatment comparisons (Figure 4, Suppl. Table 11). In PFC 17 out of 21 modules, excluding the Grey module, were enriched for LIMMA-positive results across at least one comparison group. These generally included both CIE-Drinking and CIE-NonDrinking groups. In contrast, other brain regions generally showed few modules enriched for LIMMA-positive genes and these all involved treatment comparisons with CIE-Drinking animals, although these brain regions also did not have as many LIMMA-positive genes across multiple treatment groups (Figure 4).

**Table 2:** Module names and sizes for each brain region. Module size shown in number of probesets. Module names are arbitrary colors assigned by WGCNA and do not indicate similar modules across brain regions.

| Brain Region | Module Color | Module Size | Module Color | Module Size |
| --- | --- | --- | --- | --- |
| PFC<br>(n=22) | Black | 250 | Lightyellow | 40 |
|  | Blue | 1926 | Magenta | 108 |
|  | Brown | 1588 | Midnightblue | 66 |
|  | Cyan | 71 | Pink | 192 |
|  | Darkred | 33 | Purple | 98 |
|  | Green | 513 | Red | 422 |
|  | Greenyellow | 87 | Royalblue | 37 |
|  | Grey | 1828 | Salmon | 74 |
|  | Grey60 | 49 | Tan | 81 |
|  | Lightcyan | 64 | Turquoise | 1943 |
|  | Lightgreen | 43 | Yellow | 771 |
| NAC<br>(n=25) | Black | 304 | Lightgreen | 63 |
|  | Blue | 1406 | Lightyellow | 52 |
|  | Brown | 1369 | Magenta | 212 |
|  | Cyan | 103 | Midnightblue | 97 |
|  | Darkgreen | 40 | Pink | 220 |
|  | Darkgrey | 33 | Purple | 204 |
|  | Darkred | 42 | Red | 490 |
|  | Darkturquoise | 38 | Royalblue | 44 |
|  | Green | 975 | Salmon | 118 |
|  | Greenyellow | 192 | Tan | 127 |
|  | Grey | 550 | Turquoise | 2339 |
|  | Grey60 | 66 | Yellow | 1107 |
|  | Lightcyan | 96 |  |  |
| HPC<br>(n=16) | Black | 276 | Midnightblue | 45 |
|  | Blue | 1856 | Pink | 184 |
|  | Brown | 1735 | Purple | 179 |
|  | Cyan | 54 | Red | 422 |
|  | Green | 461 | Salmon | 93 |
|  | Greenyellow | 165 | Tan | 115 |
|  | Grey | 1451 | Turquoise | 2415 |
|  | Magenta | 184 | Yellow | 630 |
| BNST<br>(n=15) | Black | 387 | Pink | 239 |
|  | Blue | 1451 | Purple | 180 |
|  | Brown | 1087 | Red | 808 |
|  | Cyan | 57 | Salmon | 72 |
|  | Green | 829 | Tan | 150 |
|  | Greenyellow | 174 | Turquoise | 2653 |
|  | Grey | 901 | Yellow | 1080 |
|  | Magenta | 201 |  |  |
| CeA<br>(n=19) | Black | 251 | Magenta | 187 |
|  | Blue | 2010 | Midnightblue | 59 |
|  | Brown | 1202 | Pink | 245 |
|  | Cyan | 70 | Purple | 125 |
|  | Green | 931 | Red | 755 |
|  | Greenyellow | 96 | Salmon | 72 |
|  | Grey | 802 | Tan | 89 |
|  | Grey60 | 35 | Turquoise | 2205 |
|  | Lightcyan | 43 | Yellow | 1035 |
|  | Lightgreen | 34 |  |  |

**Figure 4:**
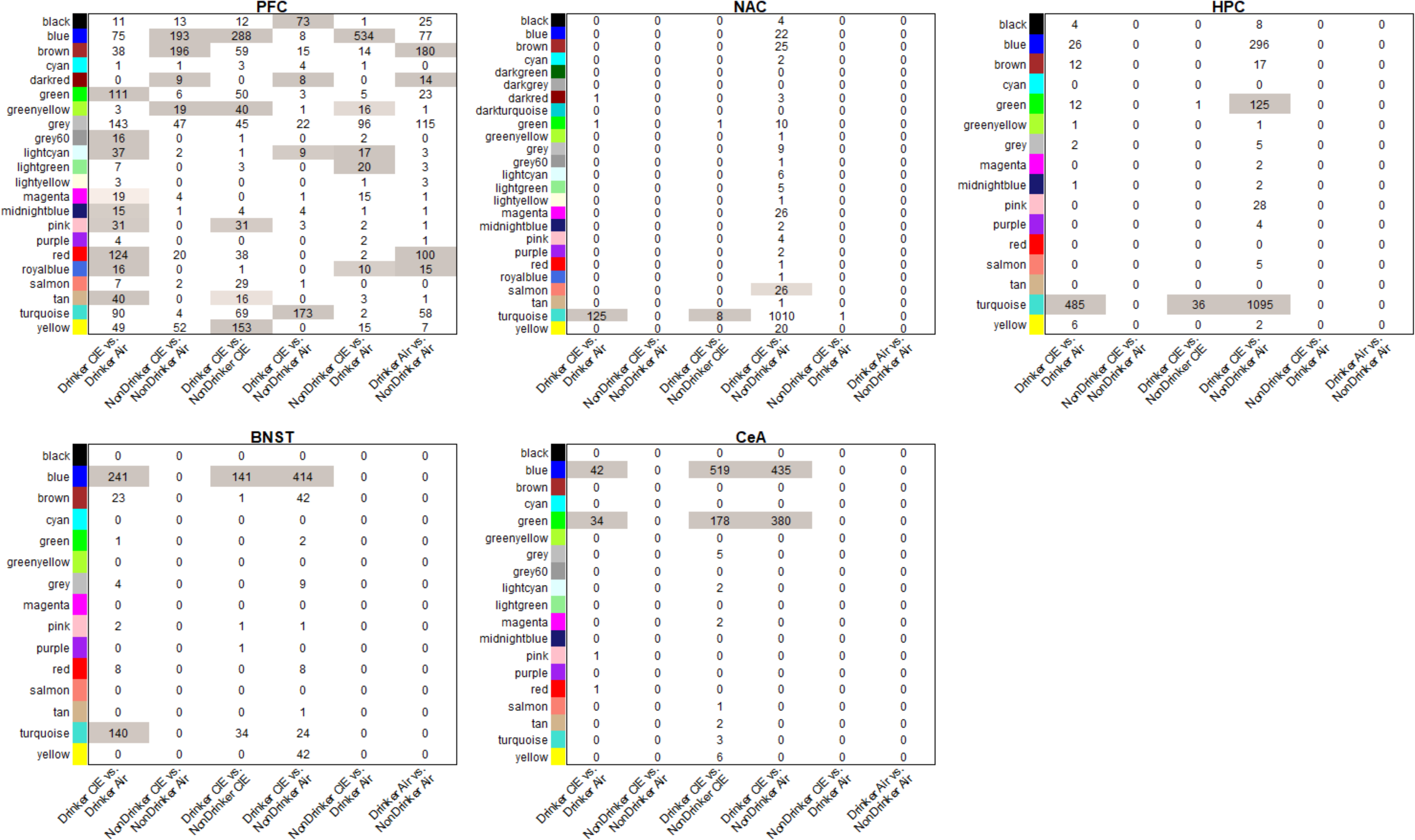
Overlap between WGCNA modules and all 6 treatment/drinking group comparisons in all brain-regions. Cell numbers indicate number of overlapping probsets between module and significantly differentially expressed genes for each comparison (LIMMA p-value ≤ 0.05). Cell color indicates significant of overlap. Significant overlap: p-value ≤ 0.05.

### WGCNA Module Phenotypic Correlations

To define WGCNA modules functionally related to ethanol drinking behaviors, we calculated Spearman correlations for module eigengenes with phenotypic data collected across the course of the experiment (Figures 5 and 8; Suppl. Figures 1-3 and Suppl. Table 5). Across brain regions, the highest correlation between drinking data and module eigengene expression was seen with ethanol intake with after CIE cycle 4, and with change in ethanol intake between baseline and CIE cycle 4. The PFC and NAC showed the largest number of modules with highly significant correlation to drinking (Figure 5 and 8). The HPC, BNST, and CeA did not show as many strong correlations to drinking, but certain modules showed module-phenotype correlations with significant p-values (≤0.05) at specific session time-points (Suppl. Figures 1-3).

**Figure 5:**
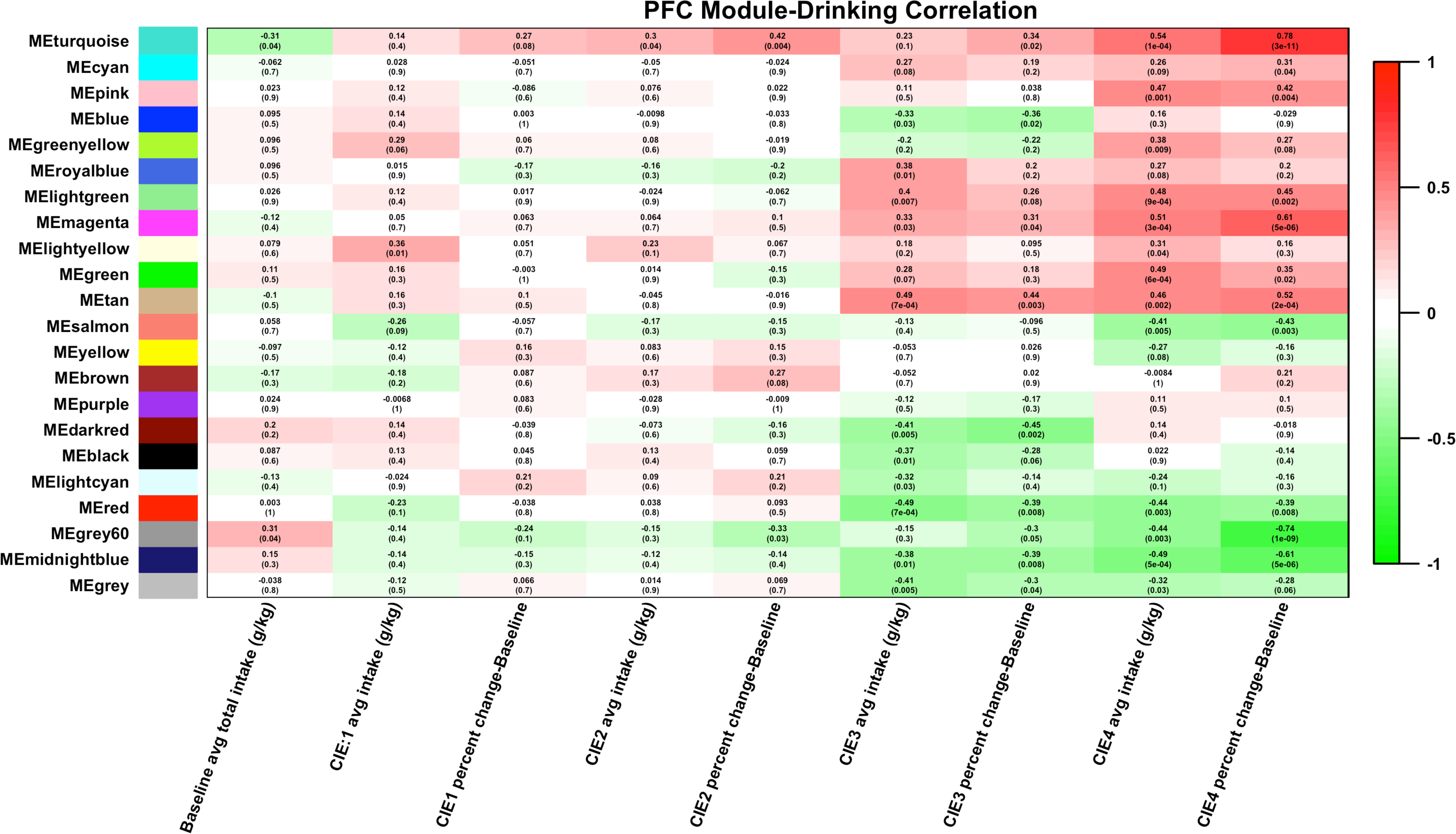
Heatmap of correlation between PFC modules and ethanol intake. Eigengene values (1^st^ principal component of gene expression) were correlated to ethanol intake measures. Cell color indicates strength of correlation (green = negative correlation, red = positive correlation).

### WGCNA Module Disruption in CIE-Drinking vs. Air-Nondrinking Groups

In addition to identifying network modules over-represented for LIMMA-positive genes or correlating with ethanol behavioral phenotypes as above, we also identified networks showing the largest degree of network structure disruption caused by CIE. Such metrics can identify more subtle changes in expression networks caused by a given treatment. For the purposes of focusing on the presumed most extreme changes, as described in Methods, we performed network disruption analysis between the Air-NonDrinking and CIE-Drinking groups. Network disruption was calculated for both the average correlation of intramodular connectivity (cor.kIM) and total connectivity (cor.kME) with full results in Suppl. Table 12. The Z_cor.kIM values gave larger numbers of disrupted modules but largely overlapped with Z_cor.kME results, and are thus discussed further here and shown in Table 3. Larger modules, in general, showed more significant disruption scores (Pearson r = −0.696, p=0.0013) but there was not a strict correspondence between the number of modules and their size vs. the number significantly disrupted by CIE treatment across brain regions. NAc showed the largest number and percentage of disrupted modules (17/24), followed by CeA (9/18), BNST (5/14), HPC (4/15) and then PFC (4/21). Thus, despite NAc only showing two modules with over-representation for Air_Nondrinking vs. CIE_Drinking regulated genes, that brain region showed the greatest percentage of modules with connectivity disrupted by CIE. This suggests a dissociation between more subtle network-level responses to CIE versus robust CIE-regulation of individual genes.

**Table 3:** Combined analysis of WGCNA module responses to ethanol. Results show modules significant for Module Disruption (Z_cor.kIM ≤ −2), correlation with Ethanol Consumption (p<0.01 for percent change vs. baseline after CIE cycle 4), or over-representation for ethanol responsive genes by LIMMA analysis of Air Nonddrinking vs. CIE Drinking (FDR<0.01). Overlap is indicated for modules present in at least two analyses (bolded module names). Module names are arbitrary colors assigned by WGCNA and do not indicate similar modules across brain regions.

| **Brain Region** | **Module Disruption by Ethanol** | **Ethanol Consumption Correlated** | **Over-represented Ethanol Regulated** | **Overlaps** |
| --- | --- | --- | --- | --- |
| PFC | Brown, Green, LightYellow, **Turquoise** | Grey60, Lightgreen, Magenta, Midnightblue, Pink, Red, Salmon, Tan, **Turquoise** | Black, Darkred, **Turquoise** | Turquoise |
| NAC | Black, Blue, Brown, Darkgreen, **Darkred**, Green, Grey60, Lightgreen, Lightyellow, Magenta, Midnightblue, Pink, Purple, **Royalblue**, Tan, **Turquoise**, **Yellow** | **Darkred**, Lightcyan, **Royalblue**, **Turquoise**, **Yellow** | Salmon, **Turquoise** | Darkred, Royalblue, Turquoise, Yellow |
| HPC | Blue, Salmon, Tan, **Turquoise** | Black, Greenyellow, Purple, **Turquoise**, Yellow | Green, **Turquoise** | Turquoise |
| BNST | Black, **Blue**, Brown, Green, Yellow | Red | **Blue** | Blue |
| CeA | Black, **Blue**, Brown, **Green**, Purple, Red, Tan, Turquoise, Yellow | **Blue** | **Blue**, **Green** | Blue, Green |

Results show modules significant for Module Disruption ( $Z_{\text{cor.kIM}} \leq -2$ ), correlation with Ethanol Consumption ( $p < 0.01$ for %change vs. baseline after CIE cycle 4), or over-representation for ethanol responsive genes by LIMMA analysis of Air\_Nondrinking vs. CIE\_Drinking ( $\text{FDR} < 0.01$ ). Overlap is indicated for modules present in at least two analyses (bolded module names). Module names are arbitrary colors assigned by WGCNA and do not indicate similar modules across brain regions. Primary data is from Suppl. Tables 5,11,13 and 14.

### Bioinformatics Analysis of WGCNA Modules

#### Prefrontal Cortex

As previously noted, the strongest correlations between ethanol intake and modules in the PFC were seen after the 4^th^ CIE cycle. The strongest correlations between WGCNA modules and all intake measures were between change from baseline drinking after CIE cycle 4 and eigengenes for the turquoise module (r=0.8, p-value = 1e-12), the magenta module (r=0.65, p-value = 6e-7), and the grey60 module (r=-0.72, p-value = 9e-9) (Figure 5). The magenta and turquoise modules showed Gene Ontology (GO) hits related to neuron development and synaptic transmission (Suppl. Table 6). Specific genes within these GO categories include *Ngfr, Ppp1r9a, Fgfr1, Sox1, Slc1a3* (turquoise module), and *Grin2b, Htt, Cacna1a, Ppp3ca, Rims1* (magenta module). All of these genes, individually, show significant correlation with change in drinking between baseline and CIE cycle 4 (Suppl. Tables 4-6). The green module also showed significant correlation to ethanol intake after CIE cycle 4, and to absolute and percent change in ethanol intake between CIE cycle 4 and baseline (r=0.49, p-value=6e-4 with ethanol intake, r=0.39, p-value=0.009 with absolute change from baseline, r=0.35, p-value=0.02 with percent change from baseline) (Suppl. Tables 4-6). This module had significant enrichment for regulation of neurotransmission as indicated by several GO categories (Figure 6). In addition, this module was significantly enriched for genes involved in neuron ensheathment by myelin (GO: 0007272, GO: 0008366, GO: 0042552). Myelin genes within this module include *Cd9, Lgi4, Cldn11, Olig2, Gjc3, Gas3st1*, and *Mbp* (Suppl. Tables 4-6). Using the myelin-related genes from the green module as an input list, GeneMANIA validated that those genes have shown co-expression, co-localization, or protein-protein interactions in previous published studies (Figure 6). The large turquoise module also showed a strong GO hit for chromatin modification (GO:0016568). Genes in the turquoise module within this category include many well-known chromatin modification genes such as *Dnmt1, Dnmt3b, Hdac8, Bcor, Crebbp, Ctcf, Bptf, Smarca5*, and *Smarcc1* [40–45] (Figure 7). The grey60 module also showed a significant GO hit for chromatin (GO:0000785). Genes within this category were *H1f0, Tcp1*, and *Klhdc3* (Suppl. Table 6). Of these genes, *Hdac8, Bcor, Crebbp, Ctcf, Bptf, Smarca5, Smarcc1, H1f0, Tcp1*, and *Klhdc3* were significantly correlated with change in baseline intake after CIE cycle 4 or with ethanol intake after CIE cycle 4 (Suppl. Tables 4-6).

**Figure 6:**
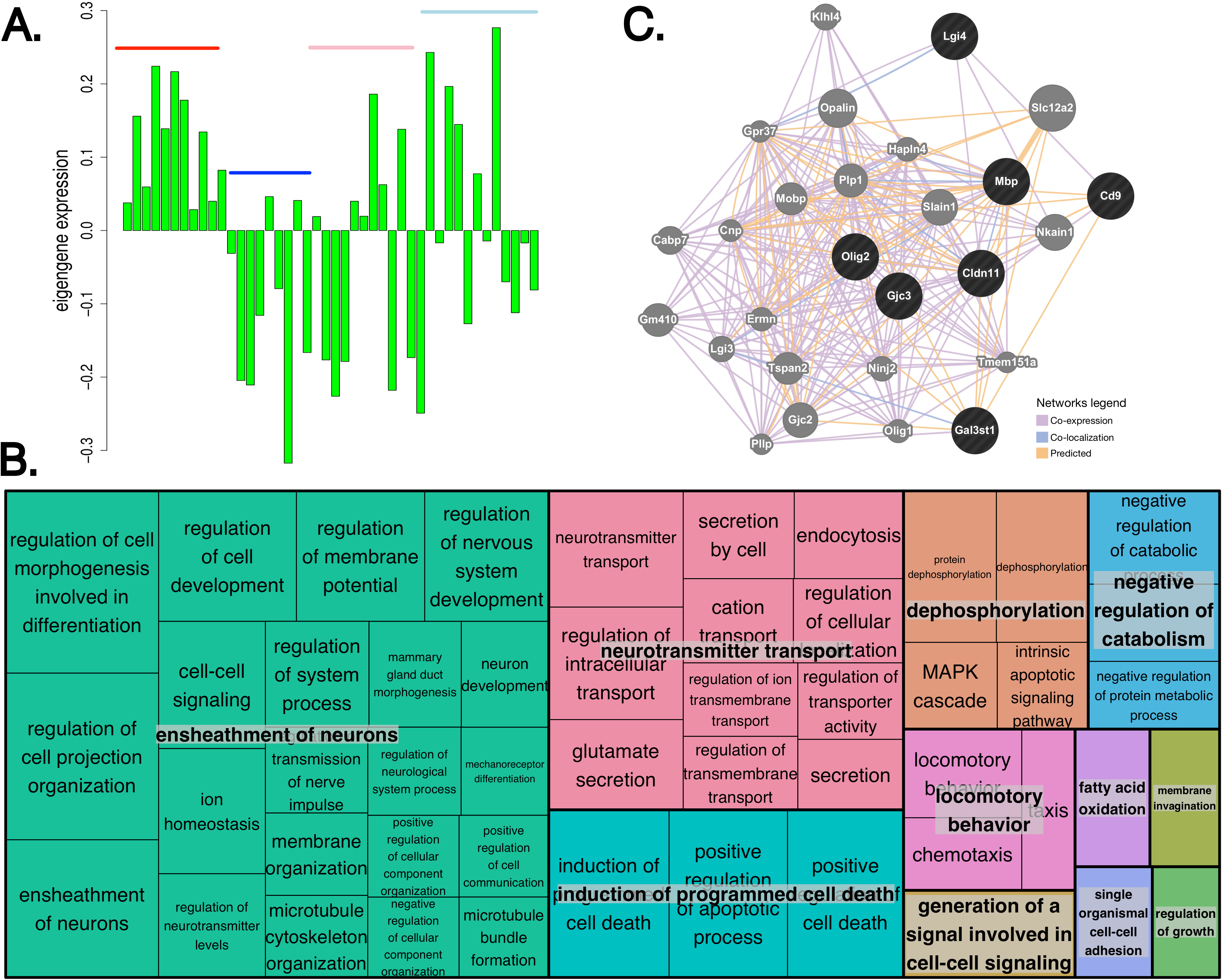
Eigengene expression for each sample in PFC green module, Gene Ontology enrichment, connectivity of myelin genes. A) Eigengene (1^st^ principal component) value from each PFC sample. Red=CIE Drinking, Blue=Air Drinking, Pink=CIE Non-Drinking, Light blue=Air Non-Drinking. B) Gene Ontology biological processes significantly enriched in the PFC green module, grouped by biological theme using REVIGO. C) GeneMANIA network generated from PFC green module genes involved in myelination.

**Figure 7:**
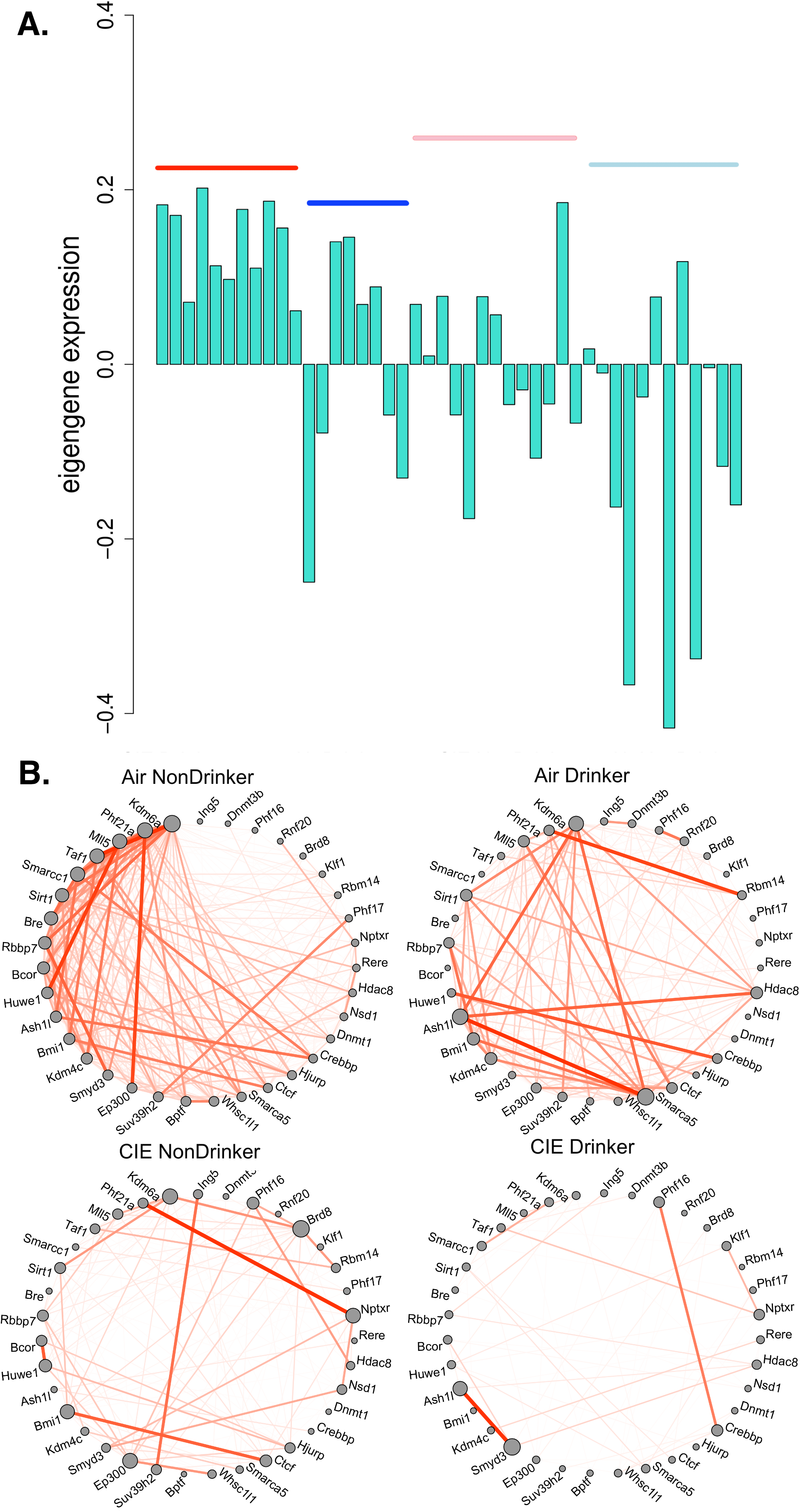
Eigengene expression for each sample in PFC turquoise module and connectivity of chromatin modification genes. A) Eigengene (1^st^ principal component) value from each PFC sample. Red=CIE Drinking, Blue=Air Drinking, Pink=CIE Non-Drinking, Light blue=Air Non-Drinking. B) Connectivity, represented by expression correlation, between genes involved in chromatin modification for each group. Line thickness and opacity represent strength of connectivity between genes.

#### Nucleus Accumbens

Patterns of module-ethanol intake correlations in the NAC were more scattered than those seen in the PFC, but the strongest correlations were still seen with intake after the 4^th^ cycle of CIE (Figure 8). These modules were the royalblue (r=0.74, p-value = 3e-10 with ethanol intake, r=0.67, p-value = 6e-8 with percent change from baseline), and salmon modules (r=-0.79, p-value = 8e-13 with ethanol intake, r=-0.47, p-value = 6e-4 with change in drinking from baseline). The royalblue module contained probesets for several subunits of the ribosomal complex (*Rps7, Rsp10, Rps13, Rps17, Rps26, Rpl12, Rpl28, Rpl32, Rpl35, Rpl36, Rpl37a, Fau*) indicating this module may play a role in regulation of protein synthesis (Suppl. Tables 5 and 7). Whereas GO hits for cellular metabolic processes, such as glucose, fumarate, glutamate, and aspartate processing, were seen in the salmon module (Suppl. Table 7). The lightyellow and yellow modules repeatedly showed significant correlation with both baseline drinking, and with drinking after each cycle of CIE. In both of these modules, however, this correlation decreased following the 4^th^ CIE cycle. Finally, several modules (blue, lightyellow, tan, magenta, salmon, and yellow) showed very strong correlation to baseline drinking. Of these, the blue, lightyellow, magenta, and yellow showed GO hits related to synaptic transmission or synaptic plasticity (Suppl. Table 7). The magenta and tan modules contained genes related to chromatin modification (Magenta: *Ing4, Ing3, Hdac1, Rbbp4, Kat5*. Tan: *Hdac9, Zbtb16*), and development (Magenta: *Rtn4, Sox9, Bmpr1b*. Tan: *Fgf9, Hdac9, Igfbp3, Zbtb16*) (Suppl. Table 7). Together, these modules indicate that, in addition to alterations in mRNA expression, CIE-induced changes in protein and metabolite populations in the NAC may be involved in the observed increase in ethanol intake (Figure 2) [19].

**Figure 8:**
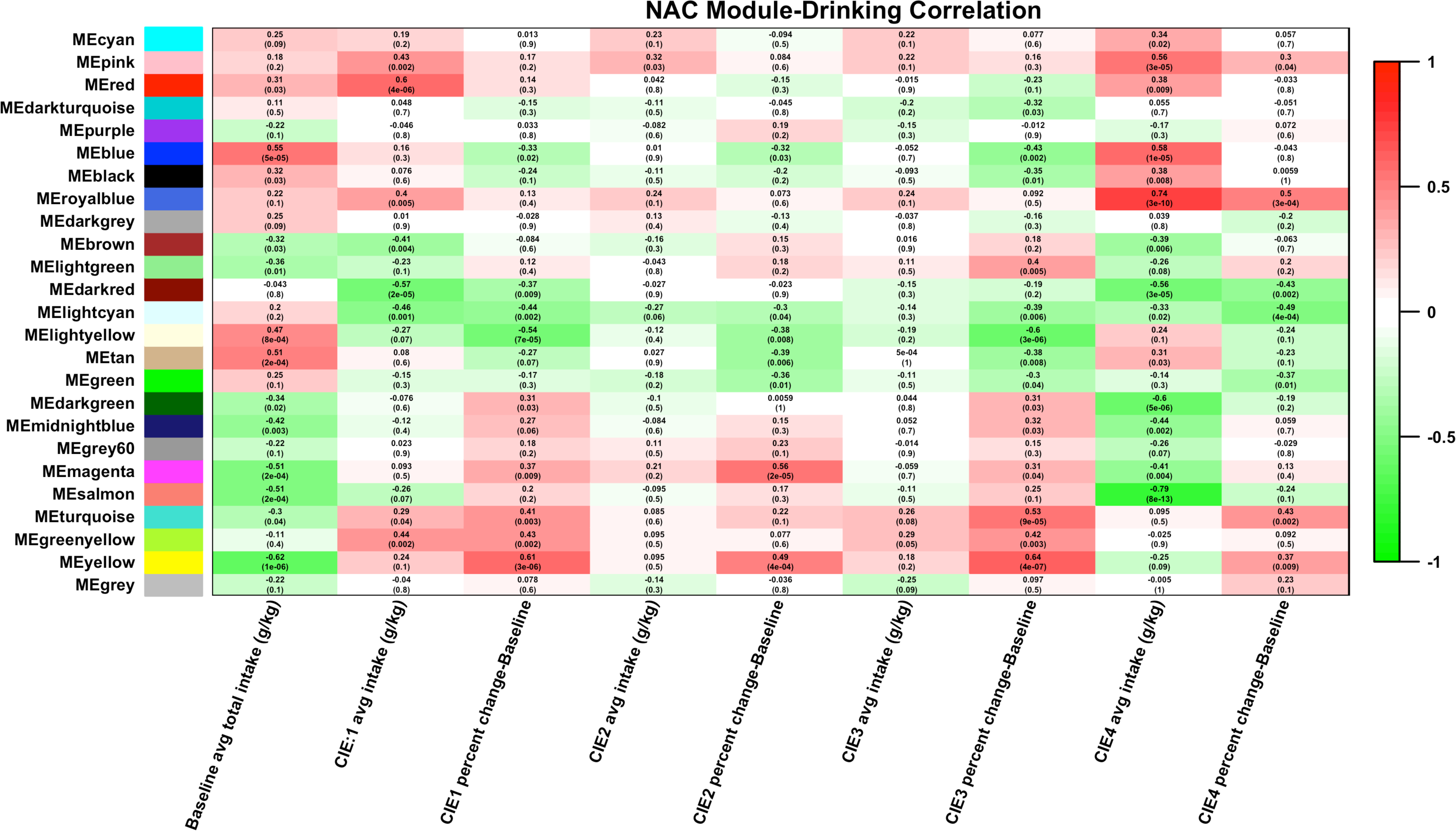
Heatmap of correlation between NAC modules and ethanol intake. Eigengene values (1^st^ principal component of gene expression) were correlated to ethanol intake measures. Cell color indicates strength of correlation (green = negative correlation, red = positive correlation).

#### Hippocampus

In the hippocampus, a noticeable pattern of module-intake correlation was also seen in after the 4^th^ cycle of CIE. In the greenyellow, black, purple, and yellow modules significant correlations were seen with change in intake from baseline to CIE cycle 4. All of these modules showed significant overlap with GO categories related to synaptic transmission (black, purple and yellow) or neuron development (purple, greenyellow, and yellow) (Suppl. Figure 1, Suppl. Table 8). The pink and magenta modules showed significant correlation to percent change in intake from baseline after CIE cycle 3 (pink module: r=0.4, p-value = 0.009, magenta module: r=0.42, p-value = 0.006). Significant correlations with intake in CIE cycle 1, and percent change from baseline intake were also seen in a few modules such as the yellow, cyan, and brown. Like the yellow module, the brown and magenta modules showed GO hits specifically for neuron development or synaptic transmission. GO analysis of the pink module showed many hits related to electron transport chain regulation, and cell motility. However, this module also showed significant overlap with two GO categories related to dendrite structure (GO:0043197, GO:0030425) (Suppl. Table 8). Genes from the pink module within these categories included *Ppp1r9a, Fbxo2*, and *Gria3. Ppp1r9a* correlated significantly with ethanol intake after CIE cycle 1 and percent change from baseline to CIE cycle 1; and *Fbxo2* and *Gria3* significantly correlated with percent change from baseline to CIE cycle 3 and CIE cycle 4 (Suppl. Figure 1).

#### Bed Nucleus of the Stria Terminalis

Fewer compelling intake correlations were seen in the BNST compared to other brain-regions. However, the turquoise and black modules showed very strong correlations to intake after the first cycle of CIE (black module: r=0.57, p-value = 4e-05 with ethanol intake, turquoise module: r=-0.55, p-value = 9e-05 with ethanol intake). Both of these modules showed multiple GO hits for synaptic transmission (Suppl. Figure 2, Suppl. Table 9). The black module also contained 4 gene ontology hits related to myelination (GO:0042552, GO:0008366, GO:0007272, GO: 0019911) (Suppl. Table 9). Genes contained within these categories included some of the known myelin building blocks such as myelin basic protein (*Mbp*), myelin-associated oligodendrocyte basic protein (*Mobp*), galactose-3-O sulfotransferase 1 (*Gal3st1*), oligodendrocyte transcription factor (*Olig2*), and Cd9 (*Cd9*) [46, 47]. Although most of these genes correlated with ethanol intake after CIE cycle 1, and with percent change from baseline to CIE cycle 1 (Suppl. Table 5), very little change in mRNA expression, with any of the 6 comparisons examined, was seen in the BNST (Suppl. Table 4). Compared to other brain-regions, the BNST also showed fewer modules with strong correlations to intake after CIE cycle 4. The red module is a notable exception, with a correlation coefficient of 0.55, and p-value of 7e-05 with change in drinking from baseline. This module also showed significant overlap with several GO categories for synaptic transmission (Suppl. Table 9). Similar to the myelin-related genes seen in the black module, however, most of the genes within these GO categories did not show significant differences in mRNA expression across treatment groups (Suppl. Table 4). In spite of these relatively level gene expression patterns, certain genes in this module did show significant correlation with ethanol intake after CIE cycle 4 and percent change in drinking from baseline to CIE cycle 4 (Suppl. Figure 2). These genes included ionotropic glutamate receptor subunits: *Gria4, Grin2b* and *Grin3a*. Metabotropic glutamate receptor 2 (*Grm2*) also correlated significantly with ethanol drinking at CIE cycle 4 and percent change from baseline.

#### Central Nucleus of the Amygdala

Module-drinking correlations seen in the CeA were sporadic, with few noticeable trends for correlation to a specific drinking measure. The two strongest correlations observed were correlations between the blue module and percent change from baseline and CIE cycle 4, and the green module with intake with CIE cycle 4 (Suppl. Figure 3). Functionally, the blue module contained several genes related to ion-mediated synaptic transmission such as *Gria4, Grin2b, Grin1, Grid2, Kcnma1, Cacnb4*, and *Cacna1a* (Suppl. Table 10). The green module, however, showed many GO hits related to chromatin modification. Several of the genes in these categories were the same as those seen in the PFC turquoise module such as *Bcor, Smarcc1, Smarca5, Bptf* and *Ctcf.* Other known chromatin remodeling genes present in the CeA green module included *Smarca4, Ncor1, Rcor1*, and *Rbbp4.* All of these genes except *Bptf*, *Rcor1*, and *Rbbp4* strongly correlated with ethanol intake after CIE cycle 4 (Suppl. Table 10). This finding is, perhaps, not surprising considering the green module as a whole (as indicated by module eigengene) also significantly correlated to ethanol drinking during the final CIE cycle (Suppl. Figure 3).

## Discussion

Through a systems biology approach we have characterized the transcriptome level response to chronic intermittent ethanol by vapor chamber with and without 2-bottle choice drinking, and identified modules of co-expressed genes in 5 regions of the mesocorticolimbic system and extended amygdala. The CIE plus drinking model has been shown; both in this study and in previous ones, to increase ethanol consumption with each successive vapor chamber cycle (Figure 2) [19, 48].

Differential expression analysis with LIMMA showed that both CIE and drinking affect gene expression in the PFC. Through overlap analysis between all comparisons of all 4 treatment groups, our results further suggested that gene expression changes in the NAC and HPC are primarily regulated by CIE, whereas in the PFC, BSNT, and CeA an interaction effect between CIE and drinking is seen (Table 1, Figure 3). Differences across treatment categories might simply reflect a linear or non-linear response to the total amount of ethanol exposure. However, the nature of the CIE and drinking model also raises the possibility that withdrawal time influences gene expression differences between the 4 treatment groups. The drinking groups, at time of sacrifice, have been abstinent from ethanol for 22 hours, whereas the non-drinking groups have been abstinent for roughly 8 days.

Network analysis with WGCNA revealed specific patterns of correlated gene expression in each brain region used in this study. This network-centric approach also allowed us to correlate both individual genes and modules of co-expressed genes directly to ethanol drinking. The strongest correlations between gene co-expression modules and drinking were seen in the PFC and NAC. These results suggest that these brain regions may have the strongest influence on the increase in drinking seen with CIE (Figure 2). The influence of the prefrontal cortex on behaviors associated with alcohol use disorders such as increased ethanol consumption and uncontrolled intake have been associated with this brain region’s role in impulse control and compulsivity [49, 50]. The nucleus accumbens, however, has been hypothesized to impact ethanol drinking behavior due to its involvement in reward [51, 52]. Therefore, ethanol-responsive gene expression changes in areas of the brain that control impulsivity and reward are implicated by network analysis in the increase in drinking seen following repeated exposure to intoxicating levels of ethanol.

One particularly striking finding was that those modules most strongly correlated with drinking after CIE exposure were consistently overrepresented for genes involved in synaptic transmission and synaptic plasticity (Suppl. Tables 6-10). This finding is not unexpected, as ethanol exposure has previously been shown to affect synaptic transmission, and synaptic architecture in several of the brain regions studied in these experiments [15, 23, 24, 53, 54]. These findings build on previous investigations into the molecular mechanisms of ethanol response in the brain, to suggest that the effect of repeated, prolonged ethanol exposure on synaptic transmission and synaptic architecture may have a direct influence on behavior both in animal models and human alcoholics. Specifically, correlated changes in expression of genes involved in synaptic remodeling in the mesocorticolimbic system and extended amygdala, in response to repeated cycles of CIE by vapor chamber, may underlie the observed increase in voluntary ethanol intake (Figure 2). In fact, recent research utilizing neuroimaging technologies have explored the effect of alcohol addiction on brain structure and function, and the relation to drinking behavior in humans [55, 56]. These studies have linked reduced grey matter volume in the medial PFC with increased risk of relapse in people with AUD [57]. SPECT and PET scanning have also shown correlations between decreased basal activity in the medial PFC during alcohol abstinence, as indicated by blood flow and glucose metabolism respectively, with poor AUD treatment outcome [58, 59]. Neuroimaging studies in mouse models are fewer; however, it is hypothesized based on previous comparative research, including those of the gene expression and behavioral response to ethanol [8, 24], that neuroplasticity changes in response to chronic ethanol exposure are highly conserved between species. Indeed, such a hypothesis has been employed in recent work using neuroimaging in rodent models to study the effect of ethanol exposure during gestation on fetal brain structure [60–62]. The results of our microarray analyses, therefore, may help shed light onto the molecular mechanisms underlying both the sustained increase in drinking observed with the CIE model, and, potentially, neuroadaptations observed in the brains of humans. Further study is needed to establish such mechanisms, and will be the topic of future research by this group.

Network analysis also identified modules in both the PFC and BNST enriched for myelin-related genes (Figure 6, Suppl. Tables 6 and 9). In the prefrontal cortex, the green module showed significant overlap with 3 GO categories related to myelination. Previous studies at our laboratory, as well as anatomical observations of the brains of human alcoholics, have suggested a role for myelination in the PFC in response to both acute and chronic ethanol exposure [13, 63-66]. Fewer studies have taken place on myelination in the BNST; however, our analyses identified the BNST black module as one with significant correlations to ethanol intake after the 1^st^ and 2^nd^ CIE cycles. Although the BNST is a lesser-studied brain region in the myelin field, this region has previously been associated with the negative reinforcing properties of alcohol [24, 67]. Our findings suggest that repeated exposures to intoxicating ethanol may also have an effect on myelination in other brain regions that have, up to this point, not been examined as often as other regions more commonly associated with ethanol related demyelination, and that changes in myelin gene expression may be another mechanism underlying increased drinking. Future avenues of study will involve examining the effect of CIE by vapor chamber on myelination in implicated brain regions, and on the effect of induced demyelination on voluntary ethanol intake with repeated exposures to prolonged levels of intoxicating ethanol.

Bioinformatic analysis also pointed to chromatin remodeling as a potential regulator of the transcriptomic response to CIE. The PFC turquoise module and CEA green module both contained genes involved in both DNA methylation [68] and members of known chromatin remodeling complexes [69–71]. *Smarcc1* has been associated with ethanol response in mouse whole brain meta-analyses [72], and *Smarca5* was found to be associated with alcohol response in network analysis of post-mortem brain tissue from human alcoholics [8]. Indeed, ethanol’s effects on epigenetic modifications to chromatin have been an area of intense study, both in humans and rodent models, during recent years [8, 73-75]. These included a study from Dr. Jennifer Wolstenholme at the Miles laboratory which found that chromatin modification genes correlated with individual variation in ethanol consumption in C57BL/6 mice [75]. Based on our findings in the other brain-regions studied, we hypothesize that this reflects the transcription level response in the brain to chronic ethanol exposure leading to downstream transcriptional regulation such as the observed changes in genes related to synaptic transmission, synaptic plasticity, and myelination.

In summary, differential gene expression and scale-free network analysis of microarray data after multiple cycles of CIE with and without intermittent access drinking has revealed brain region and treatment specific changes. Differential expression in the PFC, CEA, and BNST indicated an interaction effect between CIE and drinking; where as in the NAC and HPC, the primary effect came from CIE. Analysis of drinking patterns across multiple cycles of CIE showed that both CIE and air control mice increase their drinking, however, mice exposed to CIE drink significantly more than control. These results are in line with previous studies [19], and indicate that the CIE paradigm consistently produces progressive, lasting increases in voluntary ethanol intake in response to chronic high dose ethanol exposure. Furthermore, we have used the capabilities of network analysis through WGCNA to attempt to bridge the gap between gene expression and behavior by identifying co-expressed networks of genes in each brain region, and then correlating those networks to ethanol drinking. This strategy revealed that the most highly drinking correlated modules were seen in the PFC and NAC. In both brain-regions, as well as those with fewer significant drinking correlations, those modules with the strongest correlations to drinking, particularly after the 4^th^ CIE cycle, were enriched for genes involved in synaptic transmission or synaptic plasticity. Modules from the PFC and BNST also indicated that changes in myelin gene expression also strongly correlate to changes in drinking. These results are of particular interest as previous studies from our group have observed significant changes in myelin gene expression with acute ethanol exposure [4]. Our results also suggest a role for chromatin remodeling, particularly in the PFC and CEA, in the gene expression response to chronic, prolonged ethanol exposure. Future studies will further explore the link between chromatin remodeling and altered synaptic transmission, possibly leading to structural changes in the brain, such as altered myelination. Such changes may be mechanistically important in the drinking behavior response to chronic intermittent ethanol exposure.

## Supplementary Figure Legends

**Supplementary Figure 1: Heatmap of correlation between HPC modules and ethanol intake.** Eigengene values (1^st^ principal component of gene expression) were correlated to ethanol intake measures. Cell color indicates strength of correlation (green = negative correlation, red = positive correlation).

**Supplementary Figure 2: Heatmap of correlation between BNST modules and ethanol intake.** Eigengene values (1^st^ principal component of gene expression) were correlated to ethanol intake measures. Cell color indicates strength of correlation (green = negative correlation, red = positive correlation).

**Supplementary Figure 3: Heatmap of correlation between CeA modules and ethanol intake.** Eigengene values (1^st^ principal component of gene expression) were correlated to ethanol intake measures. Cell color indicates strength of correlation (green = negative correlation, red = positive correlation).

## Supplementary Table Legends

**Supplementary Table 1: Statistical results for drinking comparisons.** Two-way repeated measures ANOVA comparing ethanol intake in g/kg between CIE and air control (ctrl). Significance: p-value ≤ 0.05

**Supplementary Table 2: Detailed results of linear models for microarray analysis (LIMMA).** Results include log-ratio (coefficients), t-statistics for each comparison with p-values and FDR adjusted p-values. F-statistics from one-way ANOVA, F-statistic p-values, F-statistic FDR adjusted p-values, and RMA values.

**Supplementary Table 3: Summary of results of linear models for microarray analysis (LIMMA).** Significant differential expression: LIMMA two-factor model, FDR ≤ 0.01.

**Supplementary Table 4: Connectivity statistics of WGCNA modules for each brain-region**. Connectivity measures, WGCNA module assignment, RMA values, log-ratios, and F-statistics from one-way ANOVA, module membership (gene expression correlation to module eigengene) and module membership p-values for all genes used for WGCNA.

**Supplementary Table 5: Gene-trait correlation statistics for all probesets to ethanol intake.** Spearman rank correlation and p-values for all baseline and CIE drinking measures for all genes used in WGCNA, and Spearman rank correlation and p-values in parenthesis for all WGCNA modules (modules as a whole represented by 1^st^ principal component).

**Supplementary Table 6: Significant DAVID functional annotation for each WGCNA module in the PFC**. Gene Ontology categories, overlapping genes and significance values are shown for all categories with uncorrected significance of p-value ≤ 0.05. Also shown are Revigo ordering statistics for the same GO categories.

**Supplementary Table 7: Significant DAVID functional annotation for each WGCNA module in the NAC.** Gene Ontology categories, overlapping genes and significance values are shown for all categories with uncorrected significance of p-value ≤ 0.05.

**Supplementary Table 8: Significant DAVID functional annotation for each WGCNA module in the HPC.** Gene Ontology categories, overlapping genes and significance values are shown for all categories with uncorrected significance of p-value ≤ 0.05. Also shown are Revigo ordering statistics for the same GO categories.

**Supplementary Table 9: Significant DAVID functional annotation for each WGCNA module in the BNST.** Gene Ontology categories, overlapping genes and significance values are shown for all categories with uncorrected significance of p-value ≤ 0.05.

**Supplementary Table 10: Significant DAVID functional annotation for each WGCNA module in the CeA.** Gene Ontology categories, overlapping genes and significance values are shown for all categories with uncorrected significance of p-value ≤ 0.05. Also shown are Revigo ordering statistics for the same GO categories.

**Supplementary Table 11: Topological overlap statistics and results of overlap analysis between WGCNA modules and LIMMA significant results.** Topological overlap statistics include module topological overlap, resampled topological overlap, Z-score, p-value and FDR. Overlap results feature number of overlapping probesets, p-values, and Bonferroni corrected p-values.

**Supplementary Table 12: Module disruption results for all WGCNA modules.** Module disruption results include module size in probesets, correlation of total connectivity between the CIE drinking group and the Air Non-Drinking group (mod.cor.kME), mean correlation of total connectivity between all bootstrap networks (mean.boot.cor.kME), standard deviation of correlation of total connectivity between all bootstrap networks (sd.boot.cor.kME), Z-score of correlation of total connectivity between CIE drinking group vs. Air Non-Drinking group and mean of bootstrap networks (Z_cor.kME), correlation of within module connectivity between the CIE drinking group and the Air Non-Drinking group (mod.cor.kIM), mean correlation of within module connectivity between all bootstrap networks (mean.boot.cor.kIM), standard deviation of within module connectivity between all bootstrap networks (sd.boot.cor.kIM), Z-score of correlation of within module connectivity between CIE drinking group vs. Air Non-Drinking group and mean of bootstrap networks (Z_cor.kIM).

